# Target-aware Molecule Generation for Drug Design Using a Chemical Language Model^*^

**DOI:** 10.1101/2024.01.08.574635

**Authors:** Yingce Xia, Kehan Wu, Pan Deng, Renhe Liu, Yuan Zhang, Han Guo, Yumeng Cui, Qizhi Pei, Lijun Wu, Shufang Xie, Si Chen, Xi Lu, Song Hu, Jinzhi Wu, Chi-Kin Chan, Shuo Chen, Liangliang Zhou, Nenghai Yu, Haiguang Liu, Jinjiang Guo, Tao Qin, Tie-Yan Liu

## Abstract

Generative drug design facilitates the creation of compounds effective against pathogenic target proteins. This opens up the potential to discover novel compounds within the vast chemical space and fosters the development of innovative therapeutic strategies. However, the practicality of generated molecules is often limited, as many designs focus on a narrow set of drug-related properties, failing to improve the success rate of subsequent drug discovery process. To overcome these challenges, we develop TamGen, a method that employs a GPT-like chemical language model and enables target-aware molecule generation and compound refinement. We demonstrate that the compounds generated by TamGen have improved molecular quality and viability. Additionally, we have integrated TamGen into a drug discovery pipeline and identified 7 compounds showing compelling inhibitory activity against the Tuberculosis ClpP protease, with the most effective compound exhibiting a half maximal inhibitory concentration (IC_**50**_) of **1.9** μM. Our findings underscore the practical potential and real-world applicability of generative drug design approaches, paving the way for future advancements in the field.

## 1 Introduction

Generative drug design, a promising avenue for drug discovery, aims to create novel molecules/compounds with desired pharmacological properties from scratch, without relying on existing templates or molecular frameworks [2, 3]. While conventional screening-based approaches, such as high-throughput screening, virtual screening, and emerging deep learning-based screening [4– 7] usually hunt for drug candidates from libraries with 10^4^ to 10^8^ molecules [8–10], generative drug design enables exploration of the vast chemical space, which is estimated to contain over 10^60^ feasible compounds [11]. Consequently, it holds potential to identify underexplored classes of compounds, and novel compounds that are not in any existing library. This is especially important for target proteins without hit compounds (starting point for drug design) and those having developed resistance to current drugs.

Generative modeling techniques greatly empowers drug design. In recent years, a growing number of approaches have been proposed to guide the generation of drug-like compounds given the information of target proteins [12–17], stemming from creative artificial intelligence techniques such as autoregressive models [18], generative adversarial networks (GAN) [19], variational autoen-coders (VAE) [20], and diffusion models [12]. These approaches, by exploring the chemical space conditioned on the target of interest, have demonstrated the feasibility of target-based generative drug design with deep learning. However, validations with biophysical or biochemical assays are often missing [21], as most of the generated compounds lack satisfying physiochemical properties for drug-like compounds such as synthetic accessibility. In other words, despite generating a large number of novel compounds, existing approaches struggle to demonstrate their capability to provide effective candidates that can improve the real-world drug discovery effectiveness.

We therefore propose a method named TamGen (<u>T</u>arget-<u>a</u>ware <u>m</u>olecular <u>g</u>eneration). TamGen features a GPT-like chemical language model aiming for drug-like compound generation, inspired by the success of large language models [22]. The Generative Pre-trained Transformer [23] (GPT), backbone of large language models, has demonstrated its effectiveness in generating not only text [22] but also images [24] and speech [25], as well as understanding and solving scientific problems [26]. Here, we demonstrate that a GPT-like architecture and training strategy are also effective for generating chemical compounds, as these compounds can be represented using Simplified Molecular Input Line Entry System (SMILES) [27], a sequential representation akin to text. In addition, we introduce two modules to encode target protein and compound information, which allow target-aware generation of compounds based on protein structures and compound refinement based on seeding compounds, respectively. With benchmark test, we show that TamGen not only produces compounds with higher plausibility, but also enhances the balance between pharmacological activity and synthetic accessibility.

We applied TamGen to generate compounds against tuberculosis (TB), an infectious disease caused by *Mycobacterium tuberculosis* (Mtb). TB was responsible for 1.3 million fatalities and 10.6 million new cases in 2022 [28, 29], and the rising antimicrobial resistance (AMR) in tuberculosis necessitates urgent therapeutic innovation to tackle the disease [30, 31]. We focused on Caseinolytic protease P (ClpP), an essential serine protease in bacterial protein degradation system and an emerging novel target for antibiotic development [32–35]. Using a Design-Refine-Test pipeline powered by TamGen, we discovered 7 candidate compounds showing promising potency against Mtb ClpP, with half maximal inhibitory concentrations (IC_50_) ranging from 1.88 μM to 19.9 μM. Significantly, the compounds generated by TamGen not only enrich candidate pool for further optimization, but also provide effective anchors for hit expansion and structure-activity relationship (SAR) synthesis. These findings highlight the broad applicability and considerable potential of TamGen in target-aware drug design.

## 2 Results

### 2.1 TamGen enables target-aware compound design and refinement

We implemented TamGen with three modules: (1) compound decoder, a GPT-like chemical language model and the core component of TamGen, which lays the foundation for compound generation in chemical space; (2) protein encoder, a Transformer-based model used to encode the binding pockets of target proteins; and (3) a contextual encoder for compound encoding and refinement.

The compound decoder was pre-trained on 10 million SMILES randomly sampled from PubChem. The compound decoder adopts the autoregressive pre-training objective used in GPT, aiming to predict the next SMILES token based on preceding tokens (Fig. 1a). This training strategy allows for the sequential generation of compounds in both unconditional and conditional manners, depending on whether target information is provided or not. With this pre-training strategy, TamGen is able to learn general and diverse knowledge about a multitude of compounds from chemical databases (e.g., PubChem), without requiring any additional information such as binding proteins. This strategy enhances the generation capability of the compound decoder and improves the chemical properties of the generated compounds.

**Fig. 1.**
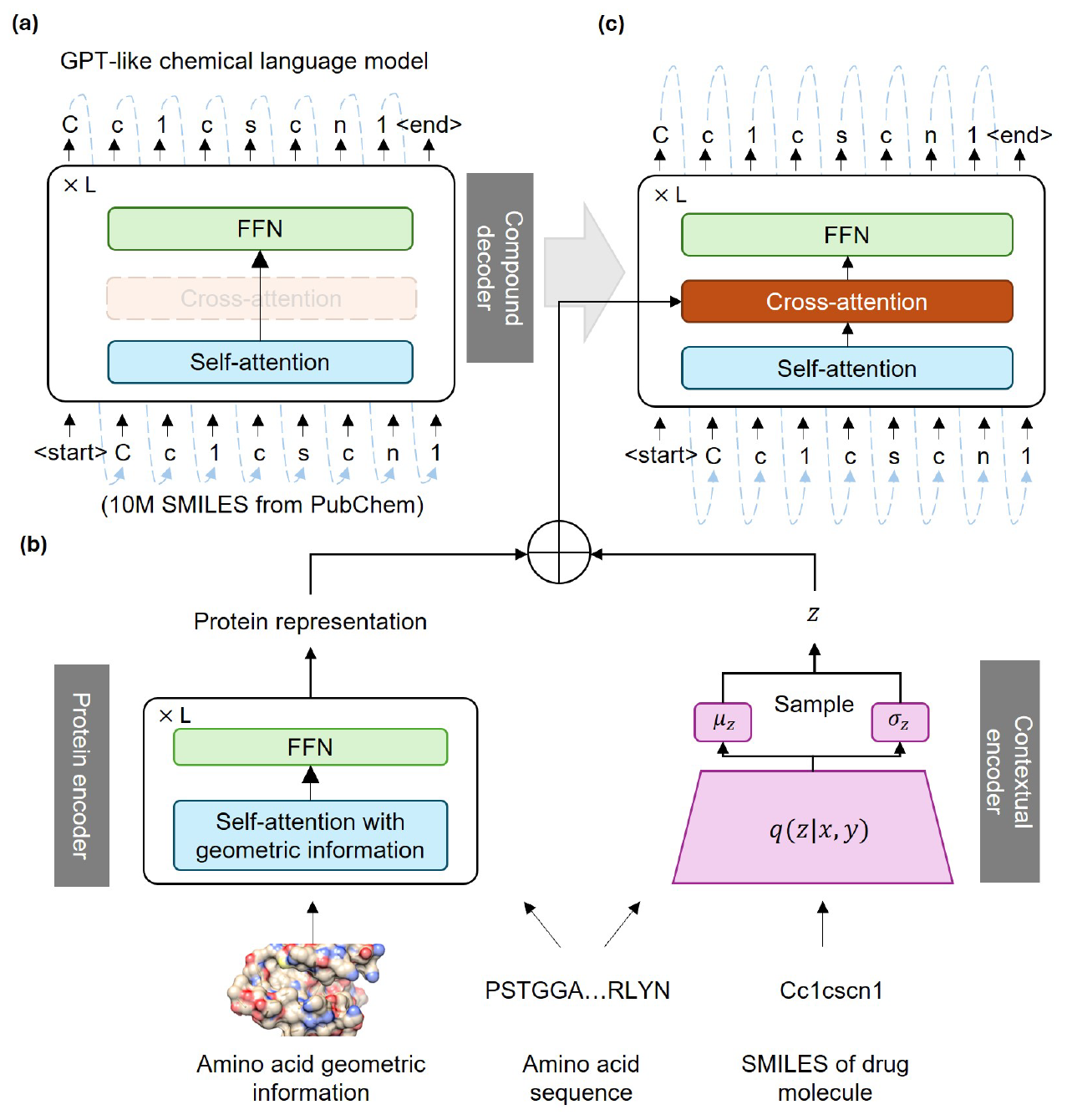
The architecture of TamGen. **(a)** The pre-training phase of the compound decoder, a GPT-like chemical language model. The model adopts standard GPT architecture, which autoregressively generates the SMILES tokens from the input. 10 million compounds randomly selected from PubChem were used for pre-training. **(b-c)** The overall framework of TamGen during the fine-tuning and inference stages. **(b)** A Transformer-based protein encoder and a VAE-based contextual encoder to facilitate target-aware drug generation and seeding molecule-based compound refinement. See Methods and Figure S1 for details. **(c)** The outputs from the protein encoder and the contextual encoder are integrated and forwarded to the compound decoder via a cross-attention module.

The protein encoder was developed to comprehend target protein information and to facilitate the generation of drug-like compounds in a target-aware manner (Fig. 1b left). The Transformer architecture adopted by the protein encoder features a self-attention mechanism, which gathers and processes information from input sequences. Here, we designed a variant of self-attention to capture both the sequential and geometric data of target proteins (Fig. S1, see Methods for details). The protein encoder’s outputs are then directed to the compound decoder via a cross-attention module (Fig. 1c), activated only when target proteins are provided. Therefore, we are able to generate compounds from the 3D conformation of target proteins via the protein encoder-compound decoder framework.

A Variational Autoencoder (VAE)-based contextual encoder was employed to encode compounds and assist the generation process. VAEs are commonly used to create new data by learning the input data’s probability distribution and sampling from it [36]. In TamGen, the VAE-based contextual encoder determines the mean (*μ*) and standard deviation (*σ*) for any given compound **y** and protein sequence **x** pair (Fig. 1b right). Later, a vector *z* is sampled from the distribution determined by *μ* and *σ* and added to the output of protein encoder, before directed to the compound decoder (Fig. 1b right). In the training stage, the model’s objective is to recover the input compound **y**, whereas during application, the contextual encoder facilitates compound refinement once a seeding molecule is provided. The incorporation of this encoder enhances control over compound generation, enabling TamGen to be seamlessly integrated into multi-round drug optimization pipelines with human feedback. This interactive and iterative drug design capability holds the potential to increase the success rate of designed compounds and accelerate the drug discovery process.

### 2.2 TamGen is effective and efficient for generative drug design

To benchmark the overall performance of TamGen, we compared our methods against five approaches proposed recently: liGAN [37], 3D-AR [38] (there is no abbreviation for the proposed method, so we refer to it as 3D-AR), Pocket2Mol [14], ResGen [39] and TargetDiff [12]. These approaches focus on direct generation of compounds in the 3D space to match protein binding pockets with diverse deep learning techniques. Following previous practices, we evaluated these methods and TamGen on CrossDocked2020 dataset [40], a well-established benchmark dataset curated from PDBbind. CrossDocked2020 is composed of a train set with about 100,000 drug-target pairs and a test set with 100 protein binding pockets. For fair comparison with previous work, we used the same training and test data as those used in [12, 14] to fine-tune TamGen.

We generated 100 compounds for each target protein in CrossDocked2020 test set with each method respectively. Then, we evaluated the designed compounds using a comprehensive set of metrics: binding affinity to target proteins, estimated by docking scores from Autodock-Vina [41]; drug-likeness, assessed using both the Quantitative Estimate of Drug-likeness (QED) [42] and Lipinski’s Rule of Five [43] based on calculated molecular physicochemical properties; synthetic accessibility scores (SAS), estimated by RDKit as a proxy for the ease of synthesis of a compound [44]; and LogP, an indicative of molecular lipophilicity, with an optimal range of 0-5 for oral administration [45]. In addition, we quantified the ability to generate diverse compounds of each method with molecular diversity. Molecular diversity is derived from the Tanimoto similarity between Morgan fingerprints of compounds. This set of metrics provides a broad and complementary assessment of compound properties, indicating the overall efficacy of a drug design method.

While each method demonstrates strengths across certain metrics, Tam-Gen is consistently top ranked. For example, TamGen achieves either the first or the second place in 5 out of 6 metrics and exhibits the best overall performance (Fig. 2a, Fig. S2 and Table S1). This finding shows that TamGen is capable of simultaneously optimizing multiple aspects of compounds during the generation process.

**Fig. 2.**
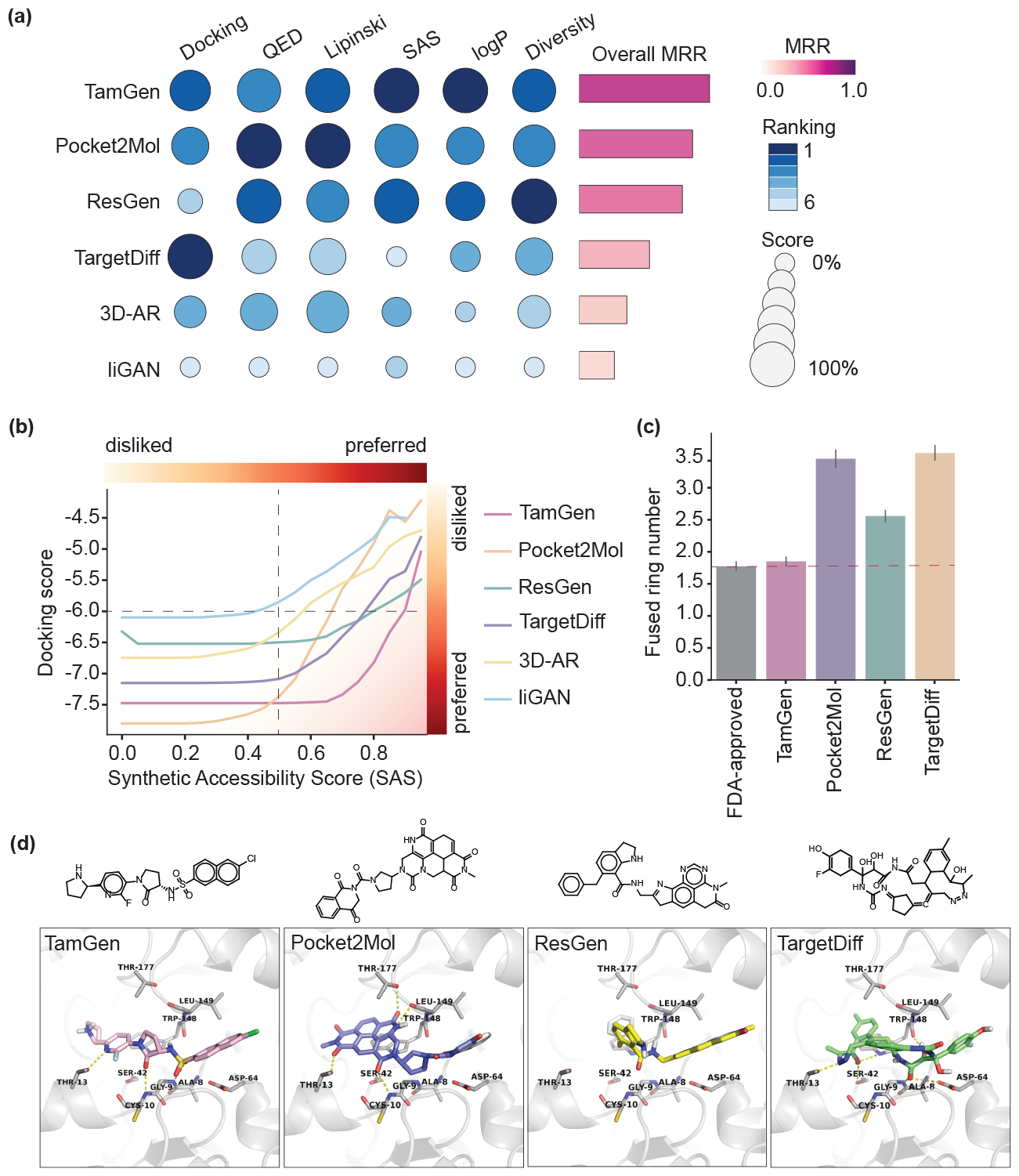
TamGen achieves the state-of-the-art performance on compound generation. **(a)** Overview of generative drug design methods ranked by overall scores for the CrossDocked2020 task. Metrics include docking score (lower scores indicate better binding affinity), quantitative estimation of drug-likeness (QED), Lipinski’s Rule of Five, Synthetic accessibility scores (SAS), LogP, and molecular diversity (Div). Scores were normalized to 0%-100% for each metric. Absolute values were used for docking score normalization. Overall scores were calculated with mean reciprocal rank (see Methods for details). See also Figure S2 and Table S1. **(b)** Average docking scores against SAS for TamGen and alternate methods. TamGen achieves more favorable docking scores for compounds with higher SAS and lower docking scores (bottom-right corner). **(c)** Barplot of the number of fused rings (see Methods for details) in FDA-approved drugs and top-ranked compounds generated by selected methods. For each method, a statistics of 1,000 compounds (100 targets *×* 10 compounds with the highest docking scores against each corresponding target) were plotted. The dashed line represents the average number of fused rings in FDA-approved drugs. Error bar, 95% confidence interval. **(d)** Example compounds generated by selected methods, and their binding poses to ClpP protein (shown as ribbons, with key residues shown as sticks).

Among the metrics, synthetic accessibility is an important factor affecting the practicality of a drug candidate, especially for novel compounds. It is worth pointing out that TamGen performs the best in terms of SAS for compounds with high binding affinity (reflected on docking scores, Fig. 2b), which are likely to possess superior bioactivity against target proteins. To discern why TamGen generates compounds with both high binding affinity and favorable SAS, we examined the top-scoring compounds generated by TamGen and other methods. Our analysis reveals that TamGen tends to produce compounds with fewer fused rings (Fig. 2c and Fig. S3). Notably, the number of fused rings in compounds generated by TamGen aligns closely with FDA-approved drugs, averaged to 1.78 (Fig. 2c and Fig. S3). Conversely, while methods involving direct 3D generation can sometimes create compounds with superior poses within binding pockets, these compounds often feature multiple fused rings (Fig. 2c-d). Prior research indicates that a higher number of fused rings may lead to lower SAS [46–48], potentially accounting for the subpar SAS scores of other methods. Moreover, a high count of fused rings is linked with increased cellular toxicity and decreased developability [48, 49]. In line with this understanding, compounds generated by TamGen display a higher similarity score to FDA-approved drugs (Fig. S4). We hypothesize that pre-training on natural compounds and employing a sequence-based generation strategy enhance the overall plausibility of compounds produced by TamGen.

TamGen also achieves the best efficiency compared to alternate methods (Fig. S5). We benchmarked the wall time to generate 100 compounds for each target of all methods using one A6000 GPU. Other methods required tens of minutes or hours to complete this task, while TamGen was able to accomplish the task in an average time of just 9 seconds. This makes TamGen 85, 154, 213 and 394 times faster than ResGen, TargetDiff, Pocket2Mol and 3D-AR, respectively.

Collectively, our results suggest that TamGen is both effective and efficient in generating novel compounds. This positions TamGen as a valuable asset for quickly identifying hit compounds for downstream development.

### 2.3 TamGen designs novel inhibitors targeting Tuberculosis ClpP protease

We next employed TamGen to design small-molecule inhibitors against ClpP. As mentioned, ClpP plays essential roles in maintaining bacterial homeostasis, rendering it a promising antibiotic target.

Apart from the previously identified Bortezomib, a peptidomimetic compound that targets the human 26S proteasome and exhibits inhibitory activity against bacterial ClpP [50, 51], there are currently no documented advanced antibiotic ClpP inhibitors. Therefore, we leverage TamGen to generate compounds targeting ClpP in *Mycobacterium tuberculosis* (Mtb), a pathogenic bacteria in urgent need for novel drug candidates.

We adopted a Design-Refine-Test pipeline driven by TamGen to identify potential ClpP inhibitors (Fig. 3). During the Design stage (Fig. 3a), utilizing the binding pocket of ClpP derived from protein structures (PDB ID 5DZK, and a ClpP-Bortezomib cocrystal structure (unpublished)), TamGen generated 2,612 unique compounds.

**Fig. 3.**
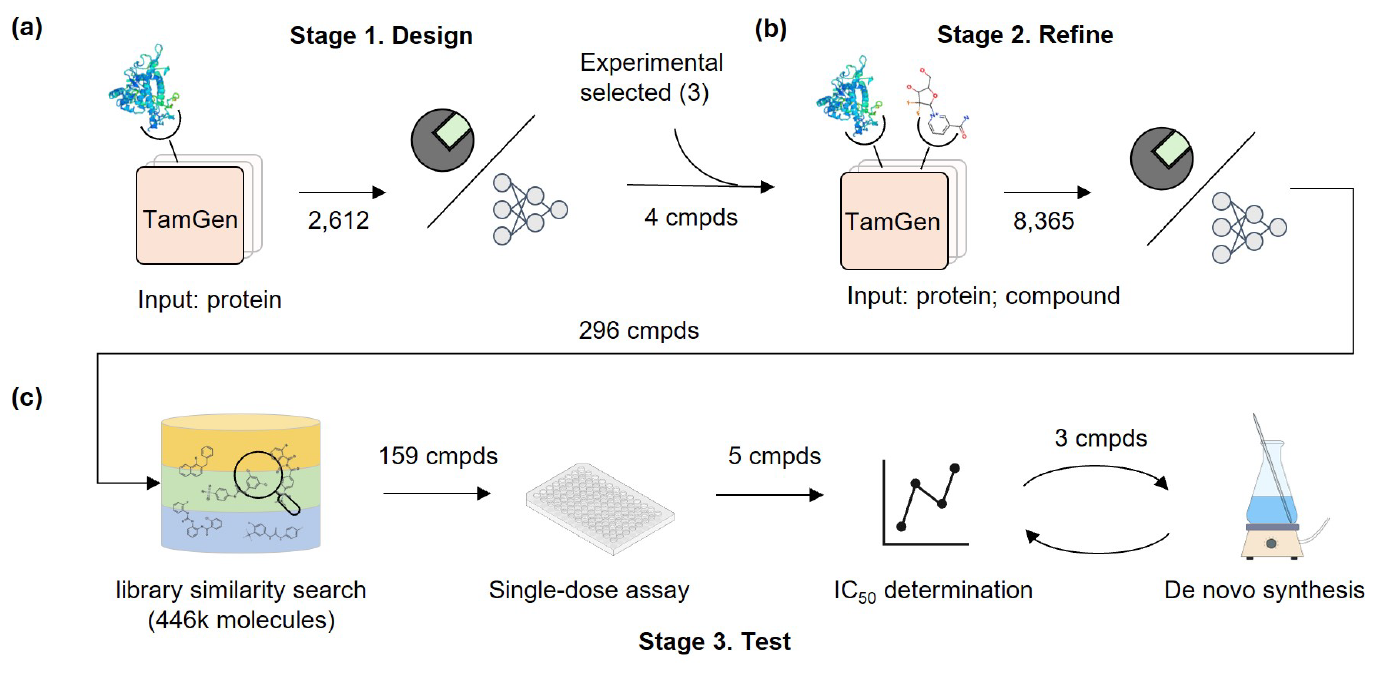
Illustration of the Design-Refine-Test pipeline for Tuberculosis drug generation. **(a)** The Design stage. **(b)** The Refine stage. **(c)** The Test stage

These compounds were then screened using molecular docking and Ligandformer, an AI model for phenotypic activity prediction [52] (see Methods for details). At this stage, we eliminated the compounds with worse docking scores compared to Bortezomib and inactive compounds predicted by Ligandformer. Peptidomimetic compounds were also excluded due to their suboptimal ADME properties (which is a known drawback of Bortezomib [53]). Finally, we identified 4 seeding compounds (green squares in Fig. 4a and Fig. S6) for the following Refine stage.

**Fig. 4.**
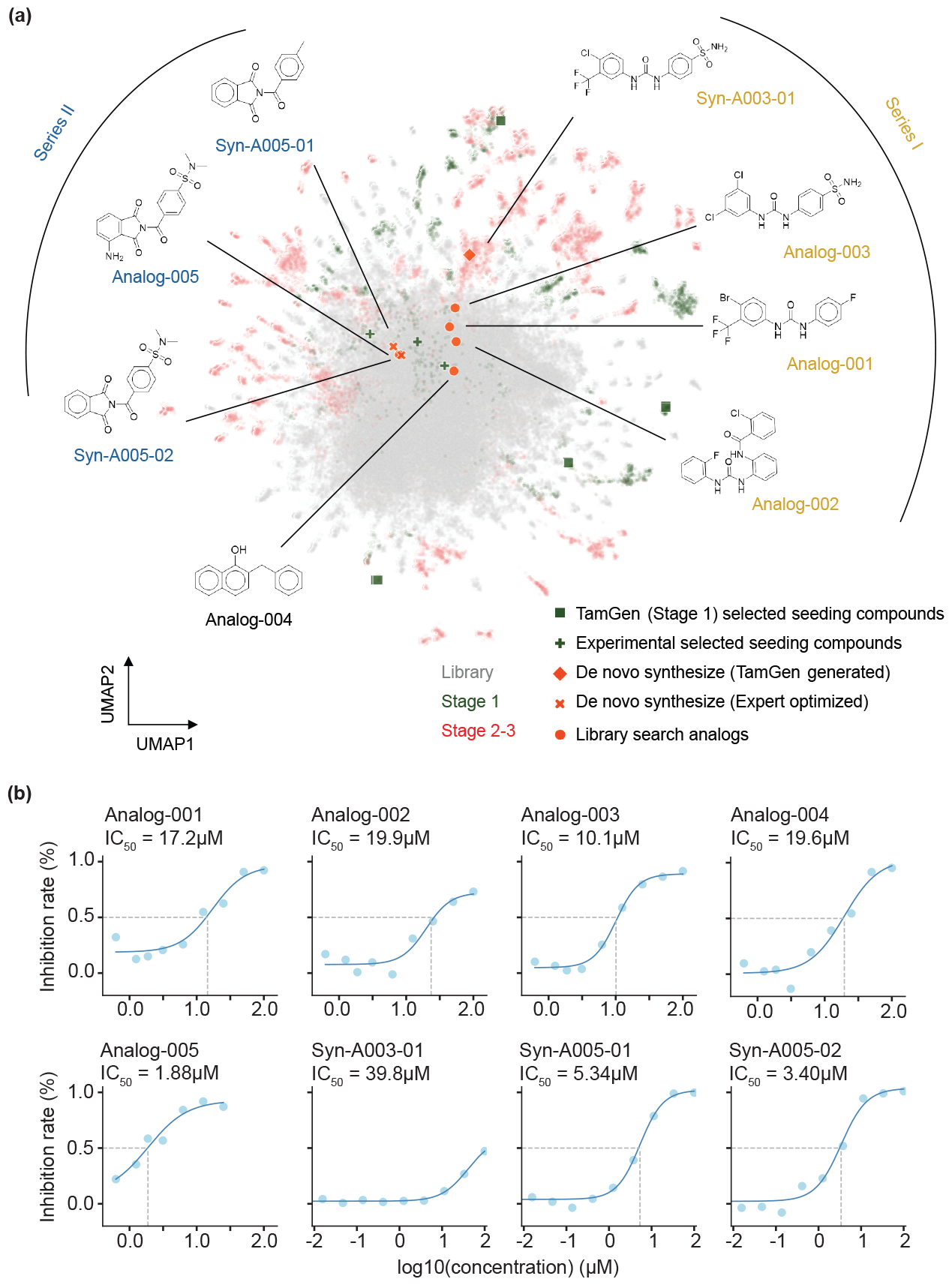
Visualization and experimental validation on designed compounds. **(a)** UMAP visualization of library compounds and key compounds identified from the Design-Refine-Test pipeline with TamGen. Gray (background): 100K compounds sampled from library. Green (background): 2,612 compounds generated at Stage 1. Red (background): 8,365 compounds generated at Stage 2. Square and plus markers in green: seeding compounds used for Stage 2 generation. Circle, cross, and diamond markers in orange red: compounds subjected to IC_50_ determinations, stratified into 3 clusters based on molecular scaffold groups. **(b)** Dose-response assays for eight compounds with DMSO as a control. See methods for details of curve fitting and IC_50_ determination.

In the Refine stage, TamGen was applied to generate compounds conditioned on both the target protein and seeding compounds (Fig. 3b). Here, in addition to the 4 representative compounds generated by TamGen, we included 3 compounds with weak inhibitory activities identified from previous experiments (IC_50_ in 100 μM - 200 μM against Mtb ClpP. Fig. S6). Conditioned on the ClpP and these 7 seeding compounds, we generated 8,635 unique compounds using TamGen, and screened the compounds following the same procedure as in the Design stage. Finally, 296 of these generated compounds were selected for the Test (biological assay) stage.

We proceeded to compare the generated compounds with molecules from existing chemical libraries. Using UMAP visualization (Fig. 4a, Methods), we observe that compounds generated by TamGen are distinguishable from those in compound libraries. This indicates that TamGen is capable of exploring untapped chemical spaces when generating potential compounds conditioned on ClpP. Moreover, the compounds generated in the Refine stage showed superior docking scores and more dispersed patterns (an indicative of molecular diversity) compared to those from the Design stage (Fig. S7). This improvement shows that a Design-Refine generation approach can effectively enhance the desired properties of the candidate pool.

### 2.4 TamGen-driven drug design yields effective inhibitors against Tuberculosis ClpP protease

To expedite the validation process and enhance the efficiency during the Test stage, we first sought commercially available compounds structurally akin to those generated by TamGen (Fig. 3c). From a 446k commercial compound library, we successfully pinpointed 159 analogues with Maximum Common Substructure (MCS) similarity scores exceeding 0.55 in comparison to any of the 296 selected TamGen compounds. Five of these analogue compounds displayed significant inhibitory effects in the ClpP1P2 peptidase activity assay, with Bortezomib serving as a positive control (Fig. S8). Subsequent doseresponse experiments revealed IC_50_ values below 20 μM for all five compounds, with Analog-005 standing out with an IC_50_ of 1.9 μM(Fig. 4b). Notably, none of these compounds have been previously documented as ClpP inhibitors, implying that TamGen may have revealed novel candidates for the treatment of Tuberculosis.

To explore the structure-activity relationship (SAR) and expand the pool of hit compounds, we synthesized three novel compounds absent from the commercial library. Considering that Analog-003 exhibited the strongest inhibitory effect in the peptidase activity assay (48% of Bortezomib, Fig. S8), we first synthesized its corresponding source compound generated by TamGen, referred to as Syn-A003-01 (Fig. 4a). Both compounds, along with Analog-001 and Analog-002, share a diphenylurea core (Series I in Fig. 4a), representing a novel scaffold for ClpP inhibitors. Interestingly, single-dose assay showed that replacing trifluoromethyl with chlorine greatly improved inhibitory activity of the compound (Fig. 4b). We reason that the replacement may have altered the charge distribution of the adjacent urine group in the compound, thereby influencing its hydrogen bonding effects. In addition, substituting sulfonamide group with fluorine also moderately improved the activity. Secondly, we synthesized two derivatives of Analog-005, the compound with the most favorable IC_50_ (Fig. 4a, SeriesII). Similar inhibition efficiency was observed in these two derivatives and Analog-005 (Fig. 4b). This result suggests a marginal contribution to the overall activity from the modified groups and provides the starting point for further modifications. Collectively, out of the eight compounds generated or inspired by TamGen, seven displayed noteworthy IC_50_ values. The high confirmation rate of TamGen-driven drug design also highlights an alternative application of generative models, specifically employing the newly generated molecules as anchors for a more effective and efficient library search. This approach allows us to alleviate the cost in screening process and surmount the challenges posed by the validation and application of novel molecule synthesis in generative methods.

### 2.5 Structural insights on the mechanisms of compound binding

To investigate the inhibitor binding mechanism, we analyzed the docking poses of two representative compounds, Syn-A003-01 (from Series I) and Analog-005 (from Series II). These two compounds were docked to ClpP structure (PDB ID: 5DZK, see Methods for details) (Fig. 5). For comparison, the binding pose of Bortezomib, derived from an unpublished cocrystal structure, was also aligned into the same crystal structure of ClpP. Similar to Bortezomib, both Analog-005 and Syn-A003-01 maintain multiple hydrogen bonding interactions with ClpP1 (a subunit of ClpP). Meanwhile, the docked pose of Analog-005 suggests that the carbonyl carbon possibly forms a covalent bond with the catalytic residue Ser98, as indicated by both the chemical mechanism and docked complex structural model. This is in accordance with the binding pose of Bortezomib, providing plausible explanation of Analog-005’s strong inhibitory activity. Interestingly, the complex structures also reveal that the sulfonamide groups of Analog-005 and Syn-A003-01 extend towards a deep pocket formed by residues Glu101, Phe102, Met150 and Asn154, a feature not observed for Bortezomib. The sulfonamide group may contribute to the binding to ClpP.

**Fig. 5.**
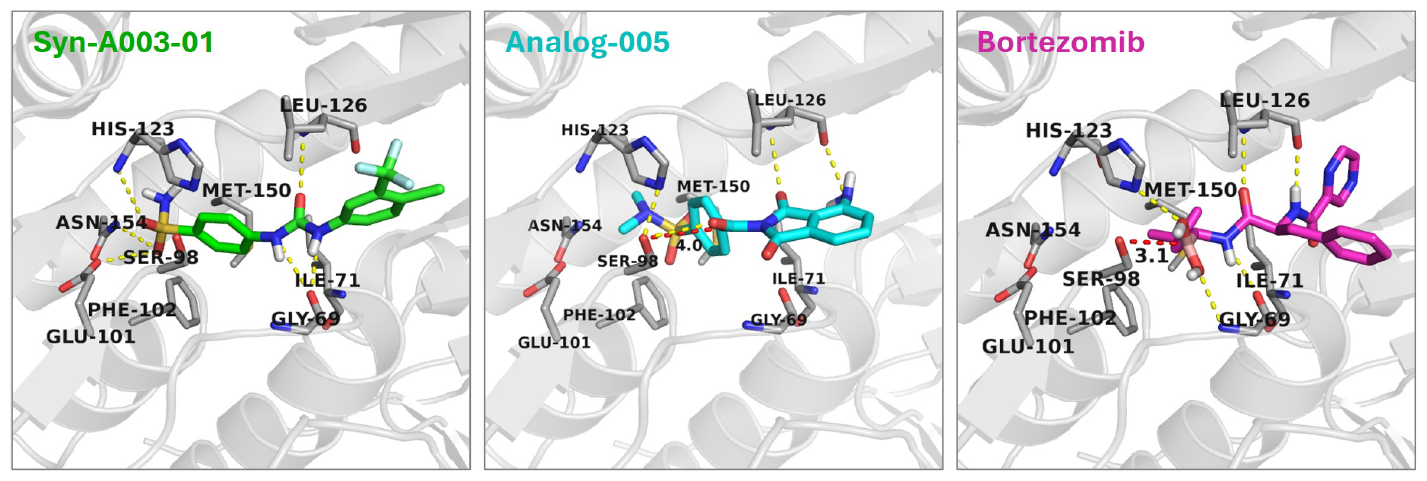
Proposed binding modes of Syn-A003-01, Analog-005, and Bortezomib against ClpP. ClpP complex 5DZK is presented in grey cartoon. Syn-A003-01, Analog-005, and the reference compound Bortezomib are shown in green, cyan, and magenta sticks, respectively. The yellow dashed lines indicate hydrogen bonds. The red dashed lines with numbers denote distances between atoms.

Altogether, through the Design-Refine-Test process powered by TamGen, we identified compounds that interact with ClpP protein in distinct modes from that of Bortezomib, thereby unveiling novel mechanisms for future ClpP inhibitor discovery. These compounds possess benzenesulfonamide and diphenylurea groups as scaffolds, which are completely different from the peptidomimetic Bortezomib, providing a possible solution to improve bioavailability and molecular stability of ClpP inhibitors. To sum up, the novelty and strong inhibitory efficacy of these compounds show potential for further development. The success of generating ClpP inhibitory compounds underscores the immense promise of TamGen in designing novel drug candidates and addressing drug-resistant Tuberculosis, implying its broad applications in drug design to treat other diseases.

## 3 Discussion and conclusions

Designing compounds that have high binding affinity to given pathogenic protein targets can speed up drug discovery process. It has been highly desirable to generate compounds based on target information and many efforts have been made to develop generative AI models to solve this challenging problem. However, few attempts have demonstrated success in real-world application. Here, we present the method, TamGen, not only achieved state-of-the-art performance in benchmark testing, but also discovered several compounds with high inhibition activities against ClpP protease of Mtb, the causative pathogen of infectious tuberculosis disease.

The success of TamGen is attributed to two major factors: (1) Chemical knowledge information embedded in the pre-trained compound decoder model, which enables the generation of high quality compounds that follow chemistry rules to possess properties for drug developments. With an ablation study, we show that pre-training is essential for producing plausible chemical compounds (Fig. S9). (2) An effective binding pocket representation that correlates to chemical compound decoding. The information of target protein binding sites is used to direct compound generation. Furthermore, TamGen can be applied to refine hit compounds reported in the literature or identified in previous rounds to generate better compounds for given targets. These designs overcome the data scarcity caused by shortage of high quality drug-target complex structures, which are usually required to learn the interactions between drug compounds and protein targets. Testing results show that TamGen is capable of generating compounds with high diversity and drug likeliness properties, increasing chances of hitting compounds that can be synthesized and further developed into drugs. This is supported by the successful design of strong inhibitor compounds against Mtb ClpP target. In the ClpP inhibitor generation case, we adopted the Design-Refine-Test workflow to iteratively improve the generated compounds. The Refine stage can be repeated multiple times by including inhibitors discovered in previous steps, so that TamGen can help further optimize the compounds and increase the chance of generating stronger inhibitors.

The pre-training of compound decoder using chemical compound information in the similar manner as GPT models is a core component of TamGen. This strategy helps overcome the data scarcity issue partially, yet, the generative AI model such as TamGen can still benefit from a larger training dataset composed of high quality target-ligand complex structures.Also, a pre-trained protein structure encoder can be applied to describe target pocket geometry information, which is currently represented using amino acid positions. Such a pre-trained model or other advanced representations for the pocket may improve generated compound qualities [54]. This is particularly important to improve the binding affinity, because the interaction information are embedded in complex structures. TamGen can be further improved to predict the compound properties, such as binding affinity, compound stability, synthesizability, and drug properties including ADME/T. As presented in this work, these properties were assessed by experts in medicinal chemistry using docking analysis and phenotypic prediction. As more 3D complex structural data along with the binding affinity or inhibition activities information become available for model training, TamGen can predict properties and rank generated compounds. Such automation will further accelerate the compound generation and facilitate experimental testing.

Generative AI models, such as TamGen, contribute to the drug discovery not only by speeding up the process, but also enable the exploration in larger chemical space beyond available compound libraries. It is expected that the information will accumulate at an accelerating pace, because the novel compounds generated by AI models will enrich the chemical knowledge once they are validated experimentally. These add-on information will in turn enhance future generative AI models. Furthermore, TamGen has demonstrated the capability of generating diverse compounds based on both binding pocket and seeding compounds. This capability enables compound refinement by providing candidates centered around the seeding compounds for follow-up research. The capability of TamGen is demonstrated in the TB drug design as an application. The same protocol can be immediately applied to design compounds for other target proteins, unleashing its power in facilitating drug discovery in general.

## 4 Methods

### 4.1 Details of TamGen

We describe the details about how to process the 3D structure input, the architectures of the protein encoder, the chemical language model, the contextual encoder and the training objective functions.

#### Preliminaries

Let **a** = (*a*_1_, *a*_2_, *· · ·*, *a*_*N*_) and **r** = (*r*_1_, *r*_2_, *r*_*N*_) denote the amino acids and their 3D coordinates of a binding pocket respectively, where *N* is the sequence length and *r*_*i*_ *∈* ℝ^3^ is the centroid of amino acid *i* (*i* is an index to label the amino acids around the binding site). *a*_*i*_ is a one-hot vector like (*· · ·*, 0, 0, 1, 0, *· · ·*), where the vector length is 20 (the number of possible amino acid types) and the only 1 locates at the position corresponding to the amino acid type. A binding pocket is denoted as **x** = (**a, r**) and [*N*] = {1, 2, *…, N* }. Let **y** = (*y*_1_, *y*_2_, *…, y*_*M*_) denote the SMILES string of the corresponding ligand/compound with a length *M* . Our goal is to learn a mapping from **x** = (**a, r**) to **y**.

#### Processing 3D input

The amino acid *a*_*i*_ *∀i ∈* [*N*] is mapped to *d*-dimensional vectors via an embedding layer *E*_*a*_. Following our previous exploration on modeling the 3D coordinates [55], the coordinate *r*_*i*_(*i ∈* [*N*]) is mapped to a *d*-dimensional vector via a linear mapping. Considering we can rotate and translate a binding pocket while its spatial semantic information should be preserved, we apply data augmentation to the coordinates. That is, in the input layer, for any *i ∈* [*N*],

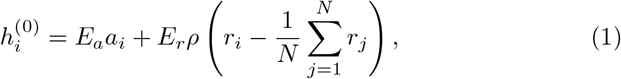

where (i) *E*_*a*_ and *E*_*r*_ are learnable matrices, and they are optimized during model training; (ii) *ρ* denotes a random roto-translation operation, and before using *ρ*, we center the coordinates to the origin. Thus we process the discrete input **x** into *N* continuous hidden representations 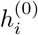 .

#### Protein encoder

The encoder stacks *L* identical blocks. The output of the *l*-th block, i.e., 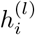, is fed into the (*l* + 1)-th layer for further processing and obtain 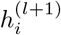 for any *i ∈* [*N*] and *l* ∈ {0} ∪ [*L −* 1]. Each block consists of an attention layer and an <monospace>FFN</monospace>layer, which is a two-layer feed-forward network as that in the original Transformer [23]. To model the spatial distances of amino acids, we propose a new type of distance-aware attention. Mathematically,

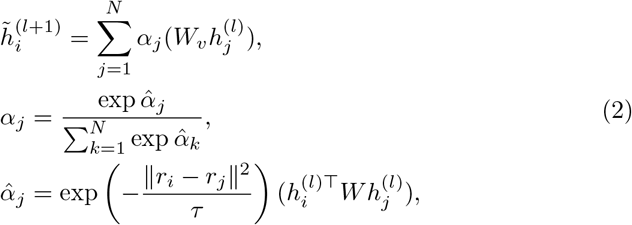

where *W* and *W*_*v*_ are parameters to be optimized, and *τ* is the temperature hyperparameter to control. After that, 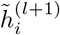 is processed by an FFN layer and obtain

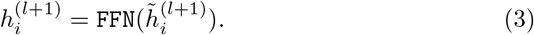

The output from the last block, i.e., 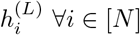, is the eventual representations of **x** from the encoder.

#### The contextual encoder

To facilitate diverse generation, we follow the VAE framework and use a random variable *z* to control the diverse generation for the same input. Given a protein binding pocket **x**, a compound **y** is sampled according to the distribution *p*(**y**|**x**, *z*; Θ). The contextual encoder (i.e., the VAE encoder) models the posterior distribution of *z* given a binding pocket **x** and the corresponding ligand **y**. The input of VAE encoder is defined as follows:

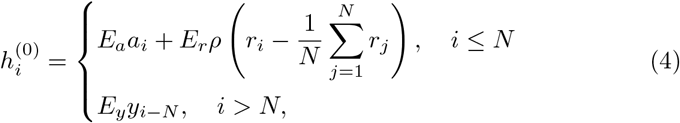

where *E*_*y*_ is the embedding of the SMILES. The VAE encoder follows the architecture of standard Transformer encoder [23], which uses the vanilla self-attention layer rather than the distance-aware version due to the non-availability of the 3D ligand information. The output from the last block, i.e., 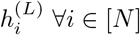, is mapped to the mean *μ*_*i*_ and covariance matrix Σ_*i*_ of position *i* via linear mapping, which can be used for constructing *q*(*z*|**x, y**), by assuming *q*(*z*|**x, y**) is Gaussian. The ligand representations, i.e., 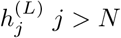, are not used to construct *q*(*z*|**x, y**).

#### Chemical language model

The chemical language model is exactly the same as that in [23], which consists of the self-attention layer and the FFN layer. We pre-train the decoder on 10*M* compounds randomly selected from PubChem (denoted as *𝒟*_0_) using the following objective function:

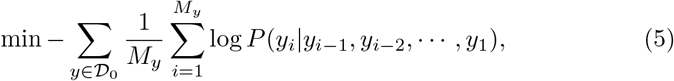

where *M*_*y*_ is the length of *y*. The chemical language model is pre-trained on eight V100 GPUs for 200k steps.

After pre-training the chemical language model, the cross-attention module is introduced to the compound decoder as shown in Fig. 1(c) (top panel). It takes all 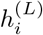 as inputs. Under the VAE variant, during training and compound refinement, the inputs are 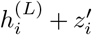, where 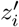 is sampled from the distribution *q*(*z*|**x, y**) introduced above. During inference, the inputs are 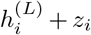 where *z*_*i*_ is randomly sampled from *𝒩* (0, *I*).

#### Training

The training objective is to minimize the following function:

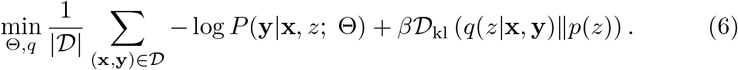

In Eqn.(6), *𝒟* is the training corpus, a collection of (pocket, SMILES) pairs; *z* in log *P* (*· · ·*) is sampled from *q*(*z*|**x, y**); *β* is a hyperparameter; *p*(*z*) denotes the standard Gaussian distribution; *𝒟*_kl_ denotes the KL divergence; Θ denotes the parameter

#### Implementation details

For the results in Section 2.2, for fair comparison with the previous methods like Pocket2Mol [14], Targetdiff [12], we use the same data as them. The data is filtered from CrossDocked [40] and there are about 100*k* target-ligand pairs. For inference, the *z* is sampled from multivariant standard Gaussian distribution. Both the pocket encoder and VAE encoder have 4 layers with hidden dimension 256. The decoder has 12 layers with hidden dimension 768. We use Adam optimizer [56] with initial learning 3 *×* 10^*−*5^. In the context of generating the compound database for Tuberculosis (TB), the current methodology incorporates an augmented dataset that includes the CrossDocked database and the Protein Data Bank (PDB), cumulatively accounting for approximately 300,000 protein-ligand pairs. To elaborate, this process involved the extraction of pocket-ligand pairs from about 72,000 PDB files. A pocket is defined on the basis of spatial proximity criteria: if any atom of an amino acid is less than 10Å away from any atom of the ligand, the corresponding amino acid is taken as part of the pocket.

### 4.2 The phenotype screening predictor Ligandformer

We utilized an adapted version of the Graph Neural Network (GNN) model as proposed in [52] to predict potential phenotypic activity. Compared with traditional GNNs, our model is designed such that the output from one layer is propagated to all subsequent layers for enhanced processing. We implemented a 5-layer architecture. Our phenotypic predictor was trained using a dataset of 18,886 samples, which are gathered from a variety of sources including ChEMBL, published datasets, and academic literature as compiled by [57]. At the inference stage, we interpreted an output value exceeding 0.69 (a threshold determined based on validation performance) as indicative of a positive sample.

### 4.3 Baselines and evaluations

#### 4.3.1 Baselines

We mainly compare our method with the following baselines:

1. 3D-AR [38], a representative deep learning baseline that uses a graph neural network to encode the 3D pocket information and direct generates the 3D conformation of candidate drugs. The atom type and coordinates are generated sequentially. 3D-AR does not explicitly generate the position of the next, by use MCMC for generation.
2. Pocket2Mol [14] is an improved version of 3D-AR, which has specific modules to predict atom type, coordinate positions and bond type.
3. ResGen [39] is also an autoregressive method of generating compounds in 3D space directly. Compared with Pocket2Mol, ResGen uses residue-level encoding while Pocket2Mol uses atomic-level encoding.
4. TargetDiff [12] utilizes diffusion models to generate compounds. Compared with the previous method, all atom types and coordinates are generated simultaneously, and iteratively refined until obtaining a stable conformation.

#### 4.3.2 TamGen without pre-training

To assess the impact of pre-training, we introduce a TamGen version without pre-training, in which the compound generator is initialized randomly. We observed overfitting when a 12-layer chemical language model was used in the non pre-trained version. Upon evaluating layers 4, 6, 8, and 12 based on their validation performance, we discovered that a model with 4 layers yielded the most optimal results.

#### 4.3.3 Mean Reciprocal Rank (MRR)

Mean Reciprocal Rank (MRR) calculation [58] is a widely used method to evaluate a method across different metrics. To elaborate, denote the rank of a method on metric *i* as *r*_*i*_. The MRR for a particular method is hence defined as 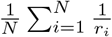, where *N* represents the total number of evaluation metrics being considered.

#### 4.3.4 Fused rings

In this work, *fused rings* denote a structural element in compounds where two or more ring structures share at least one common bond. The size of the largest group of these “fused” rings within a molecule is denoted as the number of fused rings. In Fig. 2(d), from left to right, the number of fused rings of the four compounds are 2, 5, 4 and 4 respectively.

### 4.4. Experimental details

#### 4.4.1 Peptidase activity assay

ClpP1P2 complex in Mtb can catalyse the hydrolysis of small peptides. Following previous protocols, we measure the in vitro inhibition of ClpP peptidase activity by monitoring the cleavage of fluorogenic peptide Ac-Pro-Lys-Met-AMC [59–61].

0.4 μL of candidate inhibitors, Bortezomib, or DMSO control are added into a black flat bottom 384-well plate by Echo®20 Liquid Handler and mixed with 20 μL enzyme buffer (The final ClpP1P2 dimer concentration is 50nM; reaction buffers: PIPES 30mM (pH 7.5), NaCl: 200mM and 0.005% Tween20). The solution is pre-incubated at room temperature for 2 hours. Then, 20 μL substrate buffer with Ac-Pro-Lys-Met-AMC is added (final concentration of Ac-Pro-Lys-Met-AMC is 10 μM; reaction buffer is the same with the above). Fluorescence (Ex/Em: 380/ 440 nm) is recorded for 120 min at 37^°^C.

#### 4.4.2 Single-dose response measurement

Inhibition rates of compounds were determined by Relative Fluorescence Units (RFU) compared with Bortezomib control [62, 63] and DMSO control, which is defined as follows:

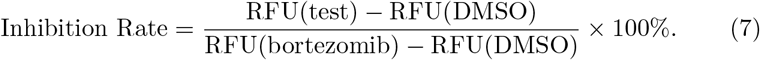

In this case, fluorescence of DMSO is seen as none inhibition (0%), and fluorescence of Bortezomib is seen as completed inhibition (100%). Compounds with inhibition rates more than 20 % at 20 μM are considered as hits.

#### 4.4.3 Dose-response assay and IC_50_ determination

To determine IC_50_, candidate inhibitors are assayed at 9 or 10 gradient concentrations. A series of candidate inhibitor, Bortezomib, or DMSO dilutions is prepared starting from a maximum concentration of 100 μM, with each subsequent concentration being half or one third of the previous one (2-fold or 3-fold dilution gradient). IC_50_ is determined by the change of recorded fluorescence (as RFU) and gradient dilution of inhibitors concentration. Non-linear fit (log(inhibitor) vs. normalized response) is used for IC_50_ curve fitting.

### 4.5 Compound generation in Design and Refine stages for ClpP

#### 4.5.1 Compound generation

Given a complex crystal structure with a protein receptor and a ligand, the center of the ligand is denoted as *c*. For each residue *i* of a protein, if its centroid *p*_*i*_ satisfies the condition ∥*c − p*_*i*_∥ *≤ τ*, i.e., within a distance cutoff *τ* from the ligand center *c*, then residue *i* is included in the pocket, where the distance cutoff *τ* is pre-defined.

In the case of ClpP complex, we first designed compounds based on published complex structure (PDB 5DZK) and our co-crystalized Bortezomib-ClpP structure. We took two values of *τ* to be 10Å and 15Å. Multiple binding sites can be extracted. We used beam search with beam size 20 to generate compounds. The *β* of the VAE was set to be 0.1 or 1. We initialized compound generation with 20 unique random seeds, ranging from 1 to 20. After removing duplicate and invalid generated compounds, we obtained 2.6k unique compounds.

During the following Refine stage, in addition to the binding pocket information, we included guiding information encoded in 4 representative compounds and 3 experimentally discovered compounds exhibiting weak inhibition activities. The parameter *τ* was set to 10Å, 12Å, and 15Å. We used beam search with beam sizes of 4, 10, and 20 for compound generation. The *β* parameter of the VAE was set to 0.1 or 1. We initiated compound generation with 100 unique random seeds, ranging from 1 to 100. After removing duplicates and invalid compounds, we obtained a total of 8.4k unique compounds.

#### 4.5.2 UMAP visualization

Compounds are converted to 1024-dimensional vectors with function `GetMorganFingerprintAsBitVect` from `rdkit` UMAP transformation [64] is performed with parameters: `n_neighbors=20, min_dist=0.7, metric=sokal michener`.

### 4.6 Ligand docking to protein target

The SMILES of generated compounds were converted to 3D structures with Open Babel program. Subsequently, AutoDock Tools was employed to add hydrogens and assign the Gasteiger charge to both the converted 3D compounds and the RCSB downloaded protein 5DZK before the docking process. The 5DZK ligand-centered maps were defined by the program AutoGrid and grid box was generated with definitions of 20 *×* 20 *×* 20 points and 1Å spacing. Molecular docking was performed with AutoDock Vina program with default settings. The predicted binding poses were visualized using the PyMol program.

## Acknowledgments

We thank Dr. Nathan Baker, Dr. Christopher M. Bishop, Dr. Sheng Ding, Dr. Marwin Segler, Dr. Ryota Tomioka and Dr. Rumin Zhang for their insightful discussions and feedback.

---

**Fig. S1.**
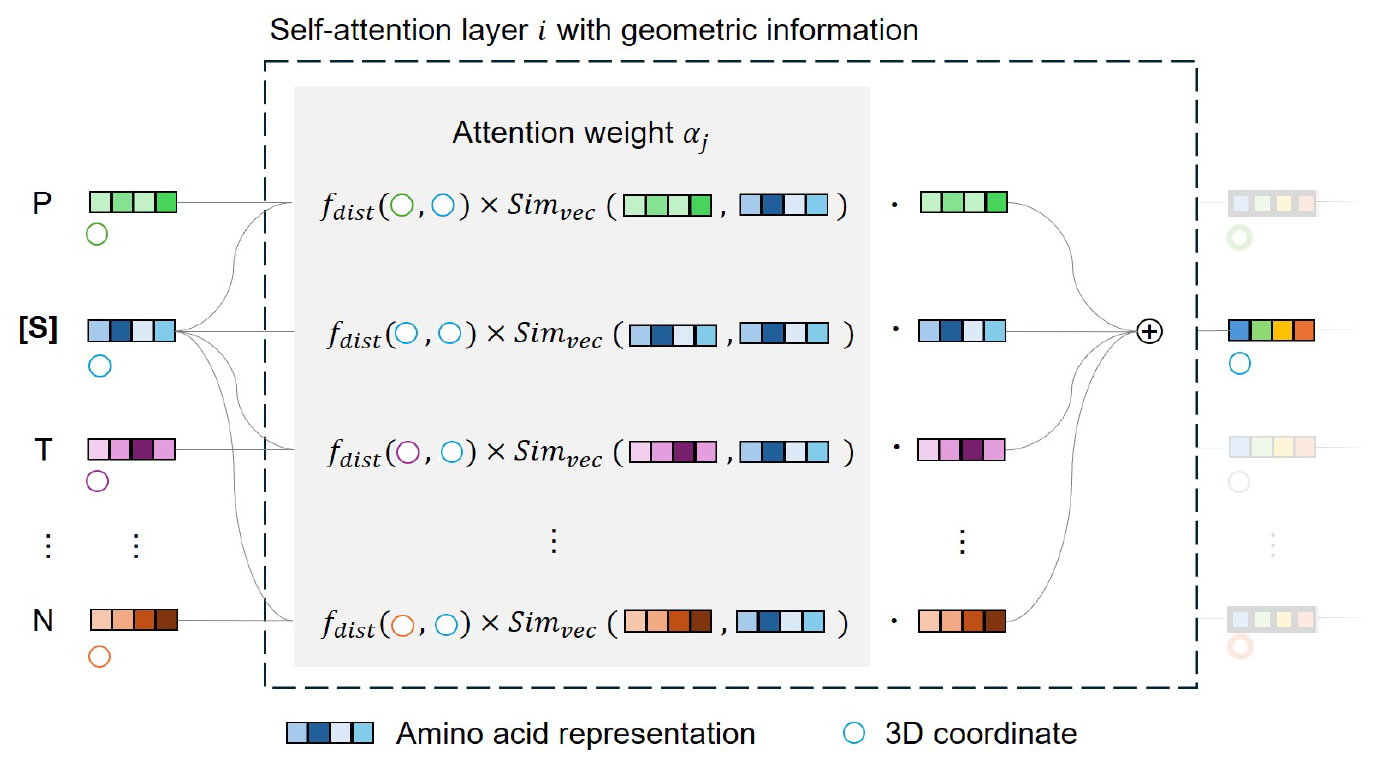
Details of the self-attention mechanism with geometric information used in the protein encoder. For each amino acid representation in layer *i*, the attention weight *α*’s is calculated as the product of the amino acid representation similarity and negative geometric distances between pairs of amino acids (i.e., exp(*−*distances^2^*/τ*) where *τ* is a hyperparameter). The output of layer *i* is then derived from the sum of the *α*’s multiplied by the amino acid representation.

**Fig. S2.**
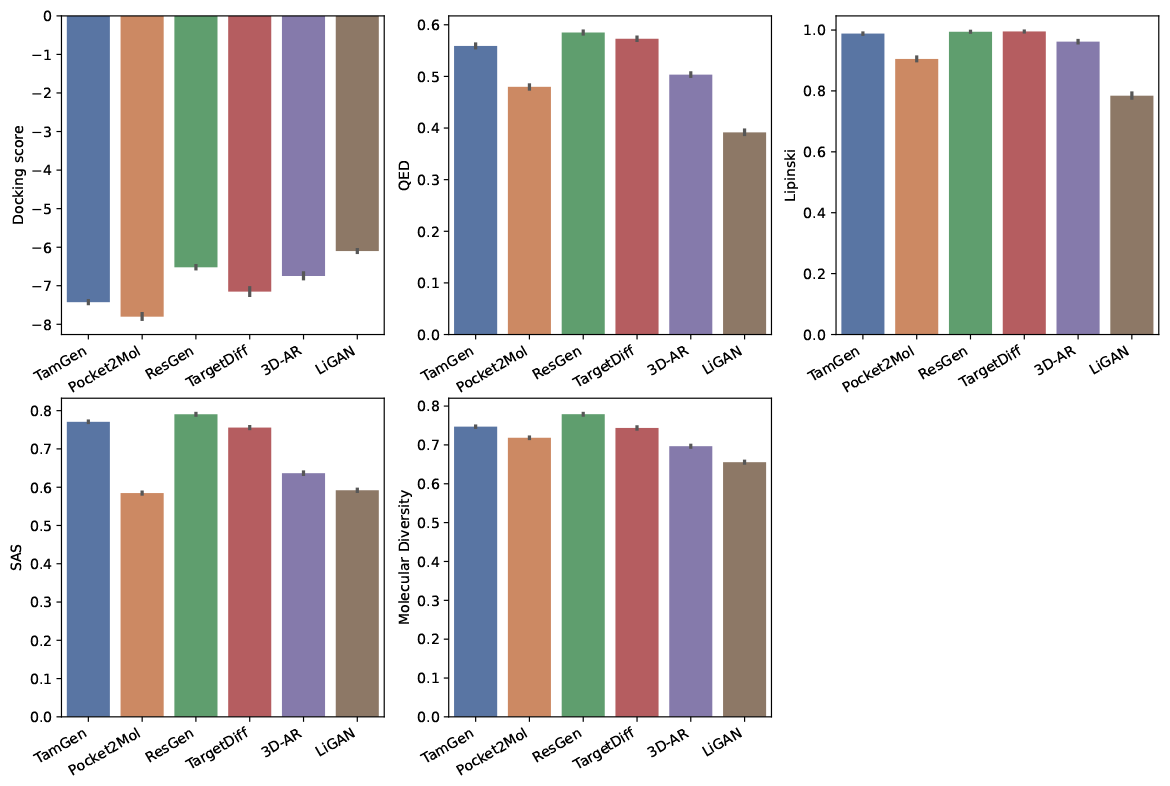
Docking scores, QED, Lipinski, SAS, and Molecular Diversity of various generative drug design methods in relation to the CrossDocked2020 task. Error bar, 95% confidence interval.

**Fig. S3.**
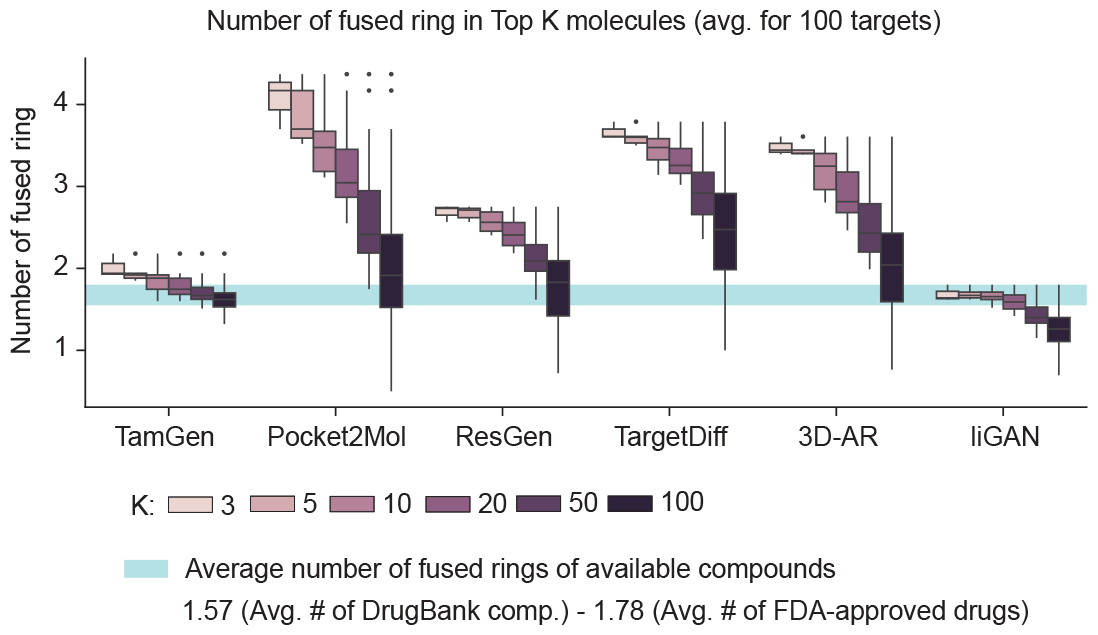
The distribution of fused ring numbers in compounds generated by different methods. *K* represents the number of compounds having top-*K* docking scores against each target protein. Center line, median; box limits, upper and lower quartiles; whiskers, 1.5x interquartile range; points, outliers.

**Fig. S4.**
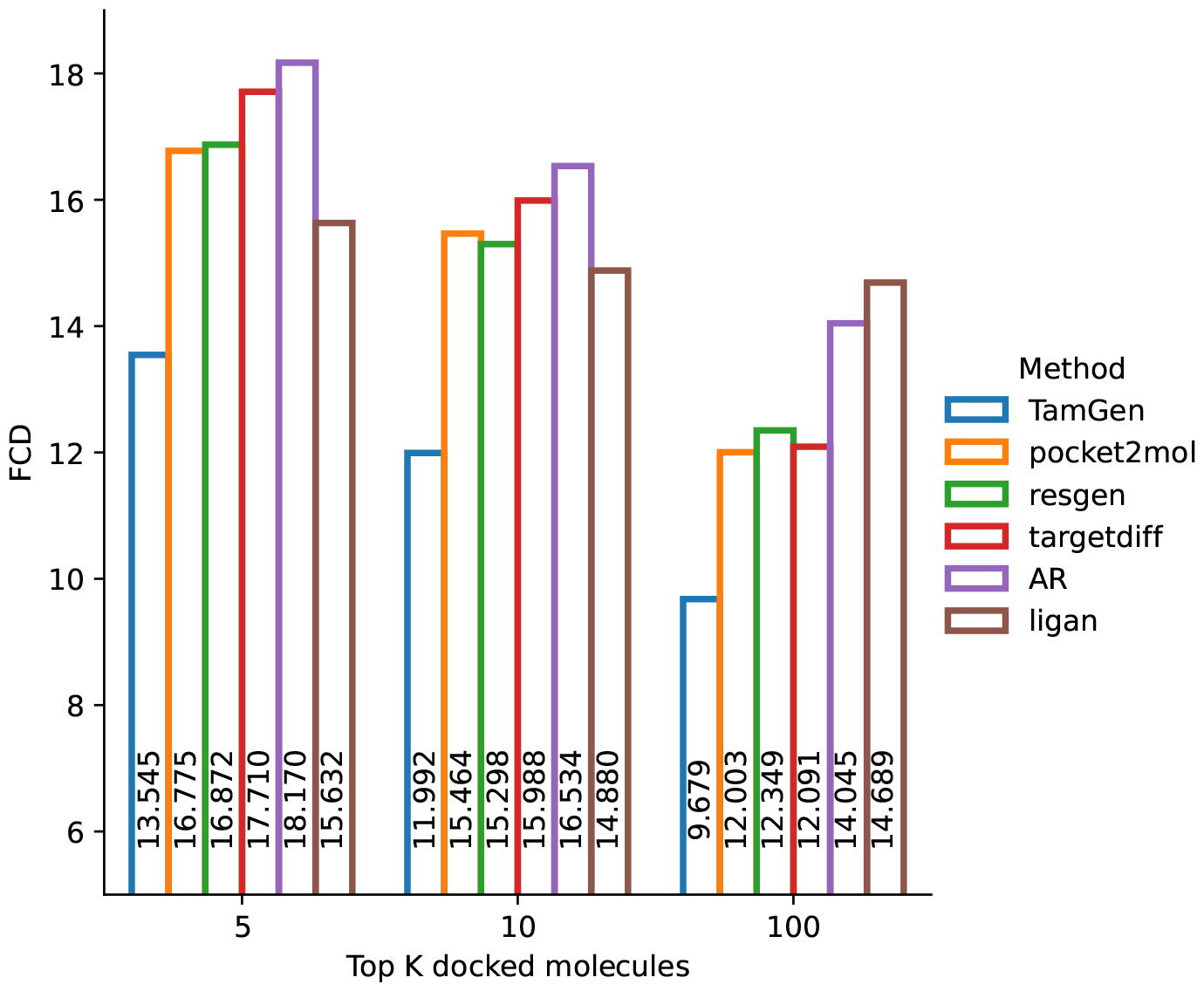
The Fréchet ChemNet Distance (FCD) similarity [65] scores between FDA-approved drugs and compounds produced by different methods. FCD is a metric that quantifies the distributional dissimilarities between two compound sets, referred to as group A and group B. In this context, group A comprises all FDA-approved drugs, while group B includes compounds generated through various methods. A lower FCD score indicates a closer distribution of the generated compounds to the FDA approved drugs, signifying their similarity. TamGen demonstrates the capability to generate compounds that are most akin to FDA-approved drugs, as evidenced by the lowest FCD scores.

**Fig. S5.**
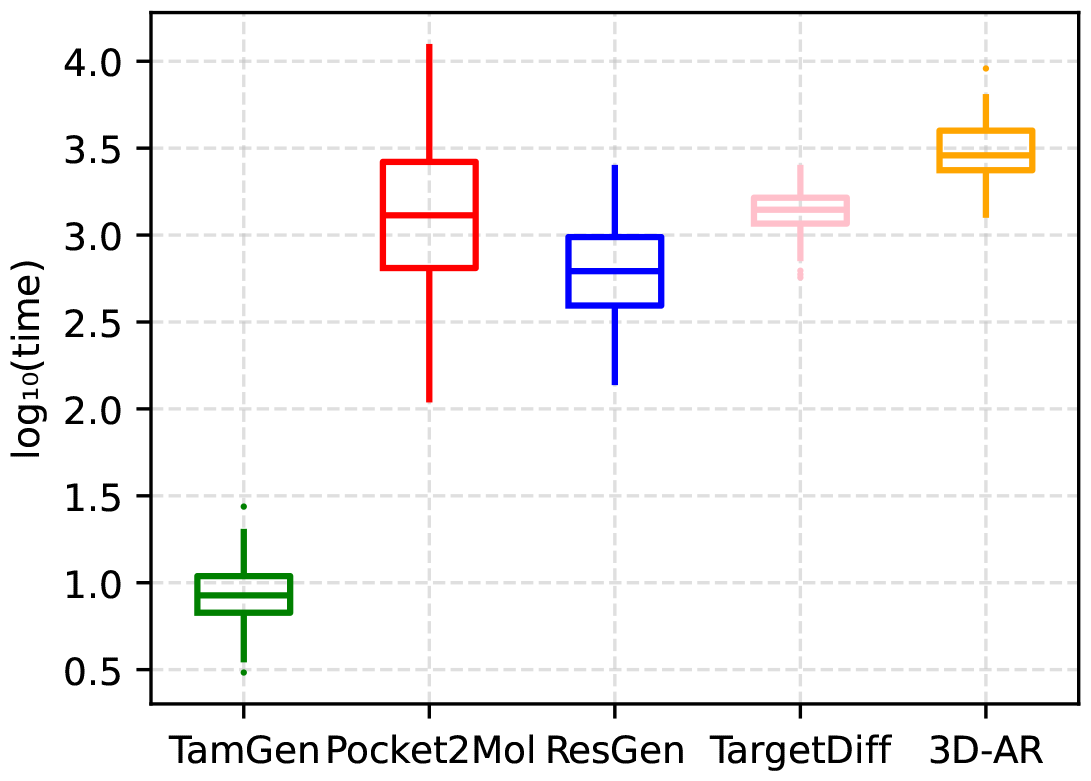
TamGen significantly outperforms alternate methods on running time. The *y*-axis is scaled using a logarithm base 10.

**Fig. S6.**
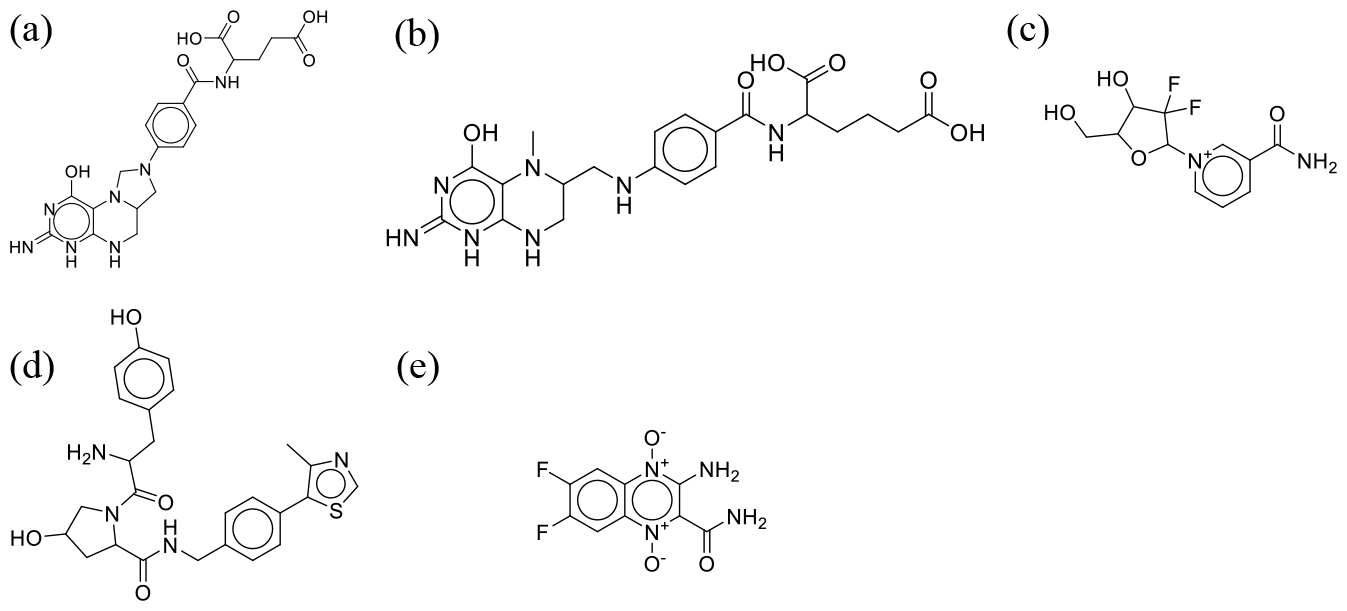
Seeding compounds for Stage 2 generation. **(a-d)** The four seeding compounds selected from the first round; **(e)**: One example of the experimental selected compound.

**Fig. S7.**
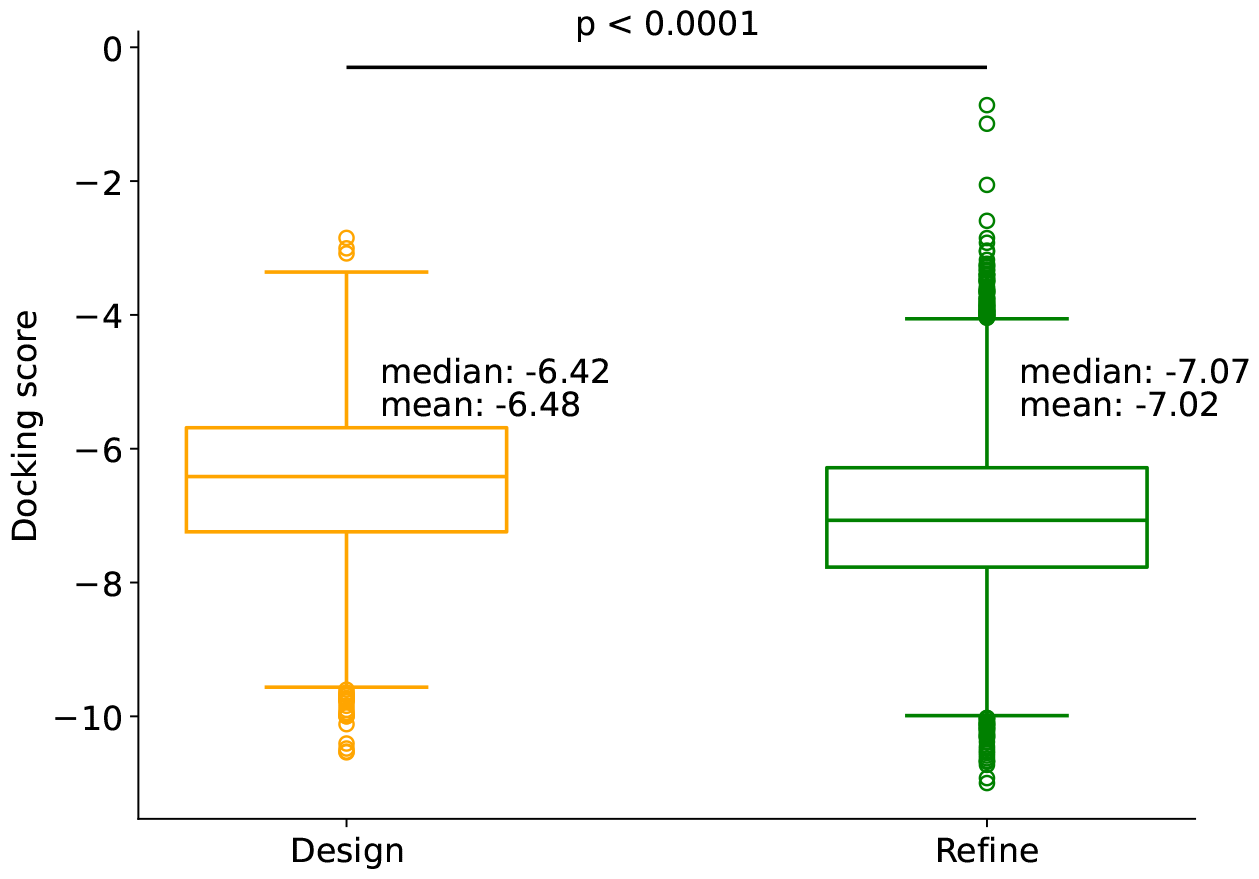
Distribution of docking scores for generated compounds against ClpP. Center line, median; box limits, upper and lower quartiles. *p*-value is calculated with Mann–Whitney U test `(scipy.stats.mannwhitneyu)`.

**Fig. S8.**
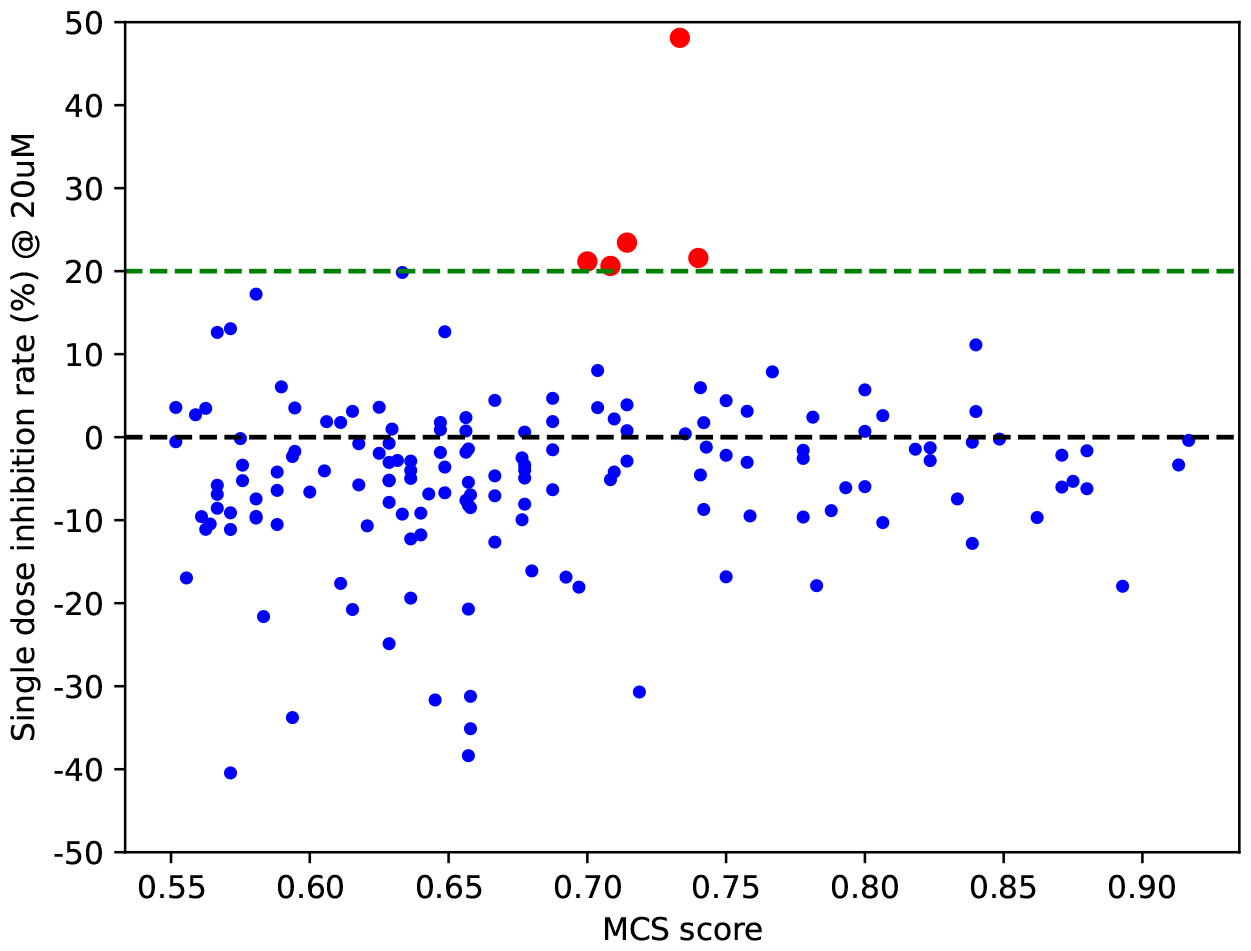
Inhibition rate of the 159 library search analogs relative to Bortezomib. All compounds were evaluated at the concentration of 20 μM. The dashed line indicates the threshold for analog selection. *x*-axis: Maximum Common Substructure (MCS) similarity scores. See Methods for details.

**Fig. S9.**
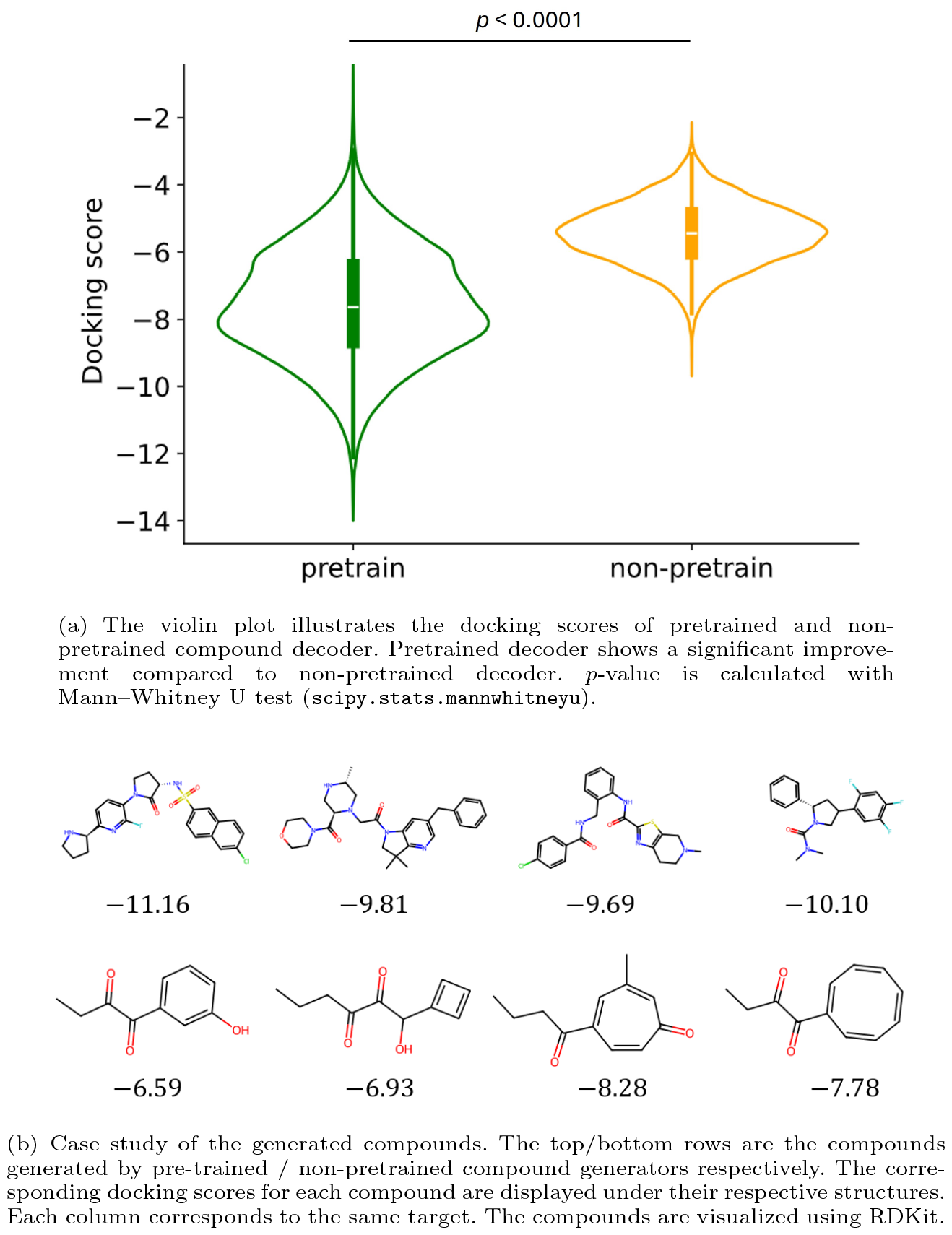
Ablation study indicates that pre-training is essential for molecule generation of the compound decoder.

**Table S1.** Compilation of performance statistics for all methods across various evaluation metrics.

| Metric \ Model | Vina Dock (↓) | QED (↑) | SAS (↑) | Diversity (↑) | LogP $\in [0, 5]$ | Lipinski (↑) |
| --- | --- | --- | --- | --- | --- | --- |
| liGAN* | -6.099 | 0.392 | 0.592 | 0.655 | 55.0% | 78.4% |
| 3D-AR | -6.746 | 0.503 | 0.637 | 0.698 | 55.0% | 96.2% |
| Pocket2Mol | -7.152 | 0.573 | 0.756 | 0.741 | 75.5% | 99.5% |
| TargetDiff | -7.802 | 0.480 | 0.585 | 0.717 | 65.2% | 90.5% |
| ResGen | -6.326 | 0.567 | 0.767 | 0.756 | 78.0% | 96.5% |
| TamGen | -7.475 | 0.559 | 0.771 | 0.747 | 87.9% | 98.8% |

**Table S2.** Resources of the analogue compounds. The index of the compounds, PubChem CID, Commercial library source and IC_50_ values are summarized.

| ID | PubChem | Commercial library source | IC<sub>50</sub> (μM) |
| --- | --- | --- | --- |
| Analog-001 | 2810424 | Maybridge Screening Collection | 17.2 |
| Analog-002 | 45503904 | Life Chemicals HTS Compound Collection | 19.9 |
| Analog-003 | 2813477 | Maybridge Screening Collection | 10.1 |
| Analog-004 | 160268 | reframeDB | 19.6 |
| Analog-005 | 4851126 | Selleck PFZ | 1.9 |

